# Tuned to Learn: An anticipatory hippocampal convergence state conducive to memory formation revealed during midbrain activation

**DOI:** 10.1101/2021.07.15.452391

**Authors:** Jia-Hou Poh, Mai-Anh T Vu, Jessica K Stanek, Abigail Hsiung, Tobias Egner, R. Alison Adcock

## Abstract

The hippocampus has been a focus of memory research since H.M’s surgery in 1953 abolished his ability to form new memories, yet its mechanistic role in memory is still debated. Here, we identify a novel, systems-level candidate memory mechanism: an anticipatory hippocampal “convergence state”, observed while awaiting valuable information, that both predicts later memory, and accounts for the relationship between midbrain activation and enhanced learning. To reveal this state, we leveraged endogenous neuromodulation associated with motivation: During fMRI, participants viewed trivia questions eliciting high or low curiosity, each followed seconds later by its answer. We reasoned that memory encoding success requires a convergence of factors, and as such, hippocampal states associated with remembered trials would be less variable than forgotten ones. Using a novel multivariate approach, we measured convergence by quantifying the typicality of spatially distributed patterns. We found that during anticipation of trivia answers, hippocampal states showed greater convergence under high than low curiosity. Crucially, convergence in the hippocampus increased with greater midbrain activation and uniquely accounted for the association between midbrain activation and subsequent memory recall. We propose that this novel convergence state in the hippocampus reflects a mechanism of its contribution to long term memory formation and that engagement of this convergence state completes the cascade from motivation to midbrain activity to memory enhancement.

## Introduction

The mysterious translation of daily life into the faulty record of memory has long compelled human wonder, and conflict. A growing literature has shown that, while some kinds of events are inherently more memorable, our ongoing motivational states are important determinants of whether and how experience is remembered (Apitz and Bunzeck, 2012; Bunzeck et al., 2009; Forkmann et al., 2013; Kennedy and Shapiro, 2009; Kentros et al., 2004; Murty and Adcock, 2014). It has been known for decades that neurotransmitters associated with motivation influence neural plasticity (Jay, 2003; Frey et al., 1990; O’Carroll and Morris, 2004; Li et al., 2003; Otmakhova and Lisman, 1996). More recent research has identified network relationships between nuclei that release these neurotransmitters and the hippocampus, long implicated in memory formation, to show that these relationships predict memory formation (Murty and Adcock, 2017; Murty and Dickerson, 2016). These studies have traced the cascade from motivational state, to engagement of neuromodulatory nuclei, to activation of hippocampal systems, concluding that neuromodulation helps the hippocampus create memories. However, they do not answer the question of *how: what computations or operations are altered to create a state conducive to memory formation*. This is perhaps unsurprising given that despite its remarkable anatomy and physiological specializations, researchers still debate the canonical function of the hippocampus and its role in memory (Basile et al., 2020; Gaffan, 1997, 2002; Graham et al., 2010; Turk-Browne, 2019)

The mesolimbic dopamine system is the major neuromodulatory system implicated in motivated learning of valuable information (Shohamy and Adcock, 2010). It has been proposed that the motivation to learn engages mesolimbic dopaminergic circuits to support plasticity in the hippocampus (Adcock et al., 2006; Gruber et al., 2014). Consistent with this account, fMRI studies in humans have shown that the motivation to learn, inspired by both intrinsic and extrinsic rewards, is accompanied by increased anticipatory activation in the dopaminergic midbrain (Adcock et al., 2006), and greater functional connectivity between the midbrain and regions in the medial temporal lobe, including the hippocampus (HPC) (Adcock et al., 2006; Gruber et al., 2014; Wittmann et al., 2007). Intriguingly, in one such study where participants were motivated by intrinsic curiosity, memory was enhanced not only for the information of interest, but also for temporally proximal irrelevant information(Gruber et al., 2014), suggesting a sustained state of enhanced encoding.

Independent lines of work suggest on the one hand that the hippocampus can express different functional states that reflect neuromodulation, and on the other that these may manifest as physiological signatures associated with distinct patterns of activity in the hippocampus. First, the hippocampus receives inputs from the dopaminergic midbrain, including ventral tegmental area (VTA) (Gasbarri et al., 1994; Swanson, 1982; Zubair et al., 2021), and midbrain projections modulate hippocampal physiology and influence performance on memory tasks (Gasbarri et al., 1996; Lisman and Grace, 2005; Lisman et al., 2011; McNamara et al., 2014; Wang and Morris, 2010). Studies of place cells in rodents have also shown that place field stability is influenced by task goals, dependent on VTA modulation (Kentros et al., 2004; Martig and Mizumori, 2011). Such shifts in response properties and circuit function have physiological signatures that could manifest in BOLD activation patterns. Indeed, in humans, dopamine receptor density has been associated with variability in the BOLD signal intensity in the hippocampus (Guitart-Masip et al., 2016). The anatomical separation of mesolimbic terminals relative to dopamine receptors in the hippocampus is ill-suited to temporally precise signals (Shohamy and Adcock, 2010) and further suggests that midbrain dopamine regulates expression of sustained functional states in the hippocampus conducive to encoding (Shohamy and Adcock, 2010; Murty et al., 2016).

Second, motivational and goal states have been shown to modulate memory-related patterns of activity in the hippocampus, detectable using multivoxel pattern analysis (MVPA) of fMRI data (Aly and Turk-Browne, 2015, 2016; Wolosin et al., 2013; Zeithamova et al., 2018). With MVPA, patterns of activity across spatially distributed voxels can be formulated as points within a high dimensional state space, with activity in each voxel constituting a single dimension. Multivoxel patterns in the hippocampus differentiate reward contexts and predict individual differences in reward-related memory benefits (Wolosin et al., 2013). When selective attention was manipulated by changing task goals, the stability of goal-related representations in the hippocampus predicted memory for the goal category (Aly and Turk-Browne, 2015). In addition to representations of contexts and goals, multivoxel patterns in the hippocampus have also been shown to differentiate processes associated with encoding or retrieval (Richter et al., 2016). Taken together, these findings suggest that variations in the hippocampal state space may reflect not only information content, but also signatures of neuromodulation.

Here, we test the premise that hippocampal state spaces reflect neuromodulation, linking hippocampal memory-related states to activation of the dopaminergic midbrain ventral tegmental area by motivation. We posit that a hippocampal state associated with successful memory formation would fall within an optimal subspace conducive for subsequent encoding. To identify the optimal subspace, we can exploit the fact that the successful formation of new memories is likely to require the convergence of multiple cognitive and physiological factors. If so, the lack of any factor could impede memory formation, yielding multiple ways to fail. An intuition for this convergence principle is captured in the opening of Leo Tolstoy’s novel Anna Karenina - *“All happy families are alike; each unhappy family is unhappy in its own way”*. The ‘Anna Karenina principle’ has been applied to the study of dynamical systems (e.g. Zaneveld et al., 2017), including the examination of human brain networks (Finn et al., 2020). Related lines of work on spontaneous brain dynamics have consistently shown better perceptual processing with reduced neural variability in sensory cortices (Arazi et al., 2019; Schurger et al., 2015). Here, we apply similar logic to identification of brain states conducive to successful memory formation. To the extent that signatures of hippocampal neuromodulation by the VTA reflect the engagement of neural states conducive to memory formation, it should be expected that they would be less variable compared to states associated with failed encoding. Thus, we hypothesized the existence of a convergence state predicting successful memory formation, when the motivation to learn is high, and reflecting midbrain neuromodulation.

In the current study, we sought to investigate relationships among motivation, midbrain activation, anticipatory hippocampal convergence states, and subsequent memory formation. To characterize convergence states, we devised a novel MVPA approach to measure trial-level variation in spatially distributed patterns. Briefly, the activation pattern for each trial is operationalized as a point in an N-dimensional state space (with N voxels). The convergence for each trial was quantified using the distance between the activation pattern and an independently defined centroid. The centroid can be thought of as a point defining the ‘average’ neural state, such that patterns closer to the centroid are considered to be more convergent than patterns further from the centroid. Higher convergence thus indicates lower variability. We applied convergence analysis to fMRI data acquired while participants engaged in a trivia quiz paradigm designed to elicit anticipatory states associated with either high or low motivation to learn, here curiosity. Replicating findings from prior work (Adcock et al., 2006; Gruber et al., 2014), we found that high motivation was associated with better subsequent recall, and with greater activation in the midbrain VTA during anticipation of answers. Using our novel analysis, we showed that anticipatory convergence in the hippocampus, but not in the medial temporal cortex, was strongly modulated by curiosity state. Higher anticipatory convergence in the hippocampus was also predictive of better subsequent recall. Critically, hippocampal convergence was strongly associated with trial-by-trial anticipatory VTA activation, and uniquely accounted for the significant association between greater VTA activation and greater subsequent memory recall. Together, our findings suggest that neuromodulation from the VTA supports memory formation by sustaining a convergence state in the HPC that is optimal for creating memories.

## Results

### Memory recall was better for high-curiosity than low-curiosity trivia

During an fMRI session, participants viewed trivia questions they had previously rated as eliciting different levels of curiosity, each followed by its answer after a variable time interval (Figure 1A). Only trivia questions that the participant indicated not knowing the answer to were included for the fMRI session (see *“Trivia question stimulus screening”* in the Methods section for details). In a memory test following the fMRI session, participants were presented with the trivia questions, and were required to recall the associated answer. Consistent with the expectation of enhanced learning in a motivated state, participants recalled more answers to trivia questions that had previously elicited higher levels of curiosity than those eliciting lower levels of curiosity (*t*(22) = 9.32, *p* < .001, *d* = 1.94, mean difference = .23, 95% CI = [.18 .28], High Curiosity: M = .63, SD = .17; Low Curiosity: M = .40, SD = .16, Figure 1B).

**Figure 1.**
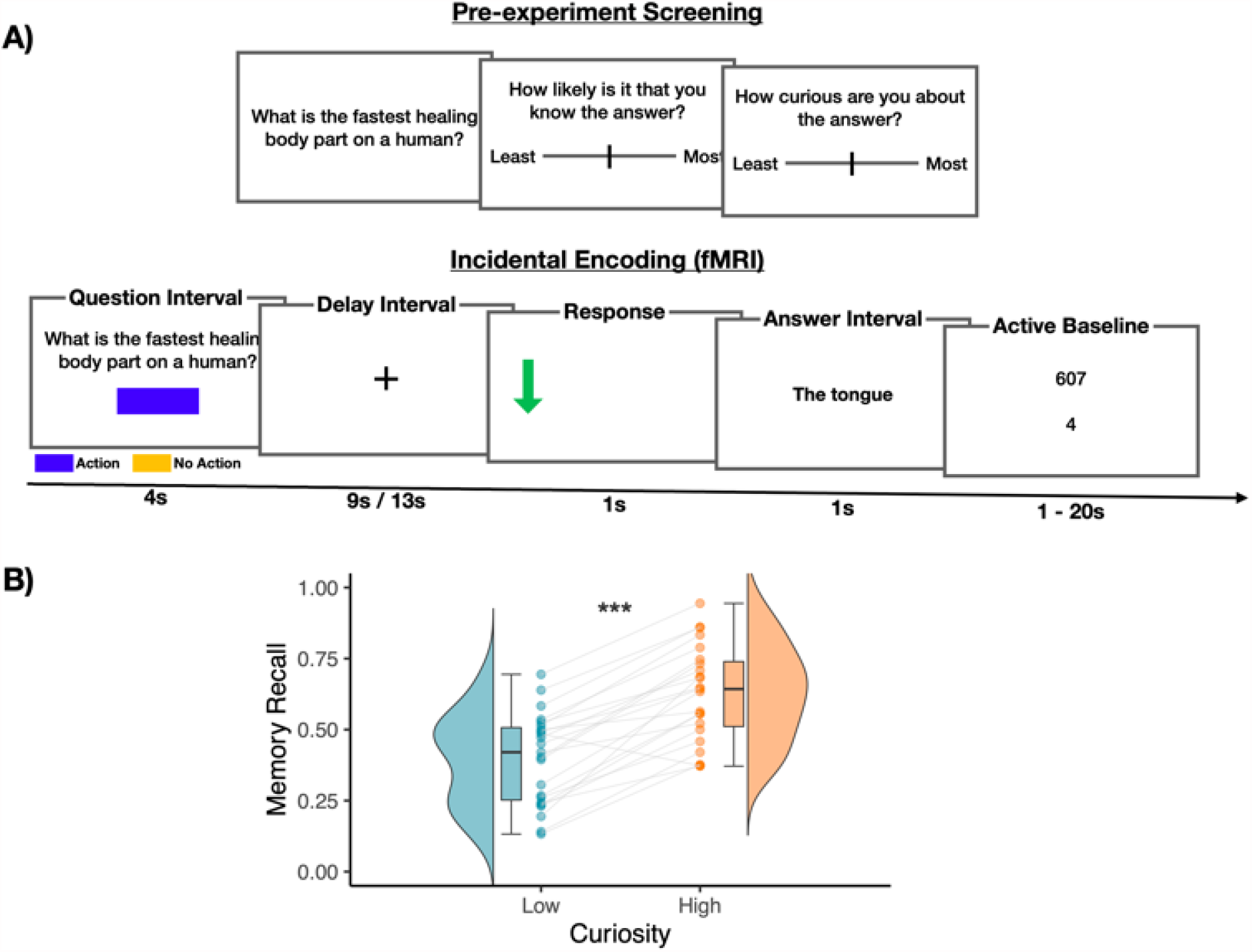
Task schematics & Memory performance. **(A)** Prior to fMRI scanning, participants were shown a series of trivia questions. For each, they were told to indicate the likelihood that they knew the answer, and how curious they were about it. Questions were excluded if participants indicated a high likelihood of knowing the answer. The included questions were separated into tertile with the 1st and 3rd tertile categorized as Low and High Curiosity questions respectively (72 questions each). During fMRI scanning, participants were shown each trivia question along with a colored rectangle that indicated the duration and action contingency of the trial. On action-contingent trials, an arrow was presented after a 9s or 13s delay. Participants indicated the direction of the arrow with a button press. This was followed by the presentation of the trivia answer. On non-action contingent trials, the trivia answer was presented immediately after the delay interval. Following the scan, participants were shown each trivia question and were required to recall its associated answer. Analyses of answer anticipation were based on activity evoked by the Question (the Question Interval). Analyses of encoding were based on activity evoked by the Answer (Answer Interval, including the response on action-contingent trials). **(B)** Box plots for memory recall performance across each condition. The upper and lower hinges of each box correspond to the first and third quartiles, while the whiskers correspond to the largest and smallest values within 1.5 times of the interquartile range. Each dot corresponds to the recall performance of each participant. ^**\*\*\***^ *p* < .001.

### VTA activation during anticipation of answers increased after high-curiosity questions and predicted better recall

We used mixed-effects models to examine if trivia questions eliciting higher curiosity also evoked greater anticipatory activation in the mesolimbic midbrain, hippocampus, and the medial temporal cortices, regions that have been associated with enhanced learning in a motivated state (Figure 2; refer to *“Univariate analysis - Effects of Curiosity on anticipatory activity”* in the Methods section for model specifications). In line with evidence of midbrain engagement during motivated learning, anticipatory activation in the VTA was greater following the presentation of high curiosity questions than following the presentation of low curiosity questions *(β* = .099, *SE* = .045, *p* = .026). In the medial temporal cortices, the perirhinal cortex showed a similar effect of curiosity *(β* = .085, *SE* = .036, *p* = .017), with greater anticipatory activation for high curiosity than low curiosity questions. While this trend was also observed in the parahippocampal cortex, it did not reach statistical significance *(β* = .08, *SE* = .042, *p* =.056). In contrast to the medial temporal cortices, anticipatory activity in the hippocampus was not significantly different between curiosity states (*β* = .019, *SE* = .040, *p* = .63).

**Figure 2.**
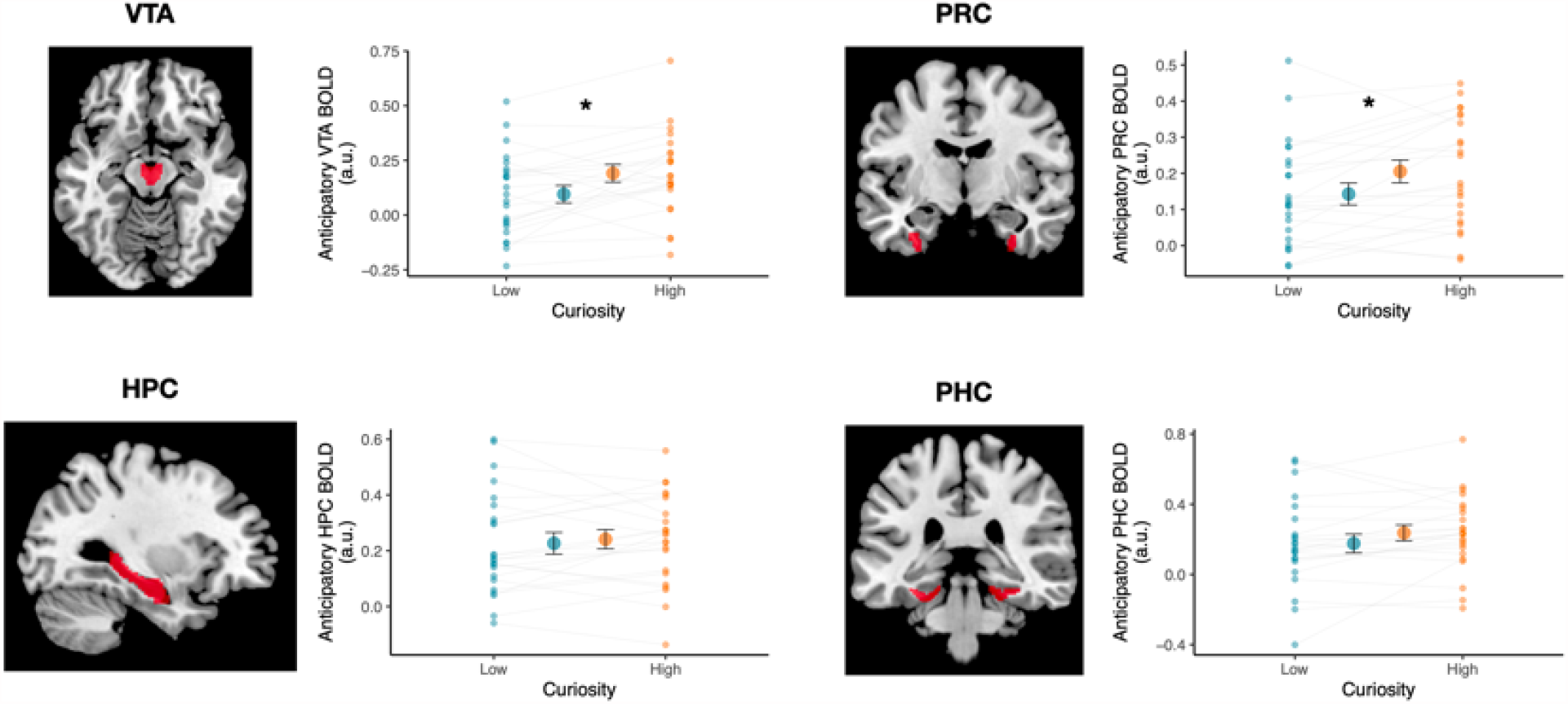
Curiosity increased univariate activation in VTA and Perirhinal cortex during anticipation of trivia answers. Anticipatory BOLD activation (i.e., during the Question interval preceding each trivia answer) was greater after High-versus Low-Curiosity trivia questions in the midbrain VTA and perirhinal cortex (PRC). Hippocampus (HPC) and parahippocampal cortex (PHC) activation did not differentiate curiosity states. Medial temporal lobe regions of interest (ROIs), included anticipatory BOLD activation (arbitrary units, a.u.) for the VTA. Red overlays on the brain images demarcate the ROIs. The large dots in each panel represent the group mean; small dots represent mean activation for each participant. Error bars represent the SEM. ^**\***^ *p* < .05.

To examine whether anticipatory activation was related to memory for subsequently presented trivia answers, we used a mixed-effects logistic regression approach with trial-level univariate activation of all ROIs included as predictors of subsequent recall (i.e. a separate regressor for each ROI). This approach allows the identification of variance that is uniquely accounted for by each of the ROIs (Figure 3). Consistent with prior findings, VTA was a significant predictor of subsequent recall, such that greater VTA activation was associated with a greater likelihood of recall (*β* = .121, *SE* = .049, *p* = .015). Univariate activation in all other ROIs was not a significant predictor of memory outcome. To ensure that this was not simply driven by greater univariate activity for the High curiosity questions, a model comparison was performed comparing model fit between a model with and without an interaction term for the Curiosity condition. The inclusion of an interaction term for VTA activation and Curiosity did not result in a better model fit (**χ**^2^ = 1.72, *p* = .190), suggesting that fluctuations in anticipatory VTA activation are related to subsequent recall performance regardless of curiosity states.

**Figure 3.**
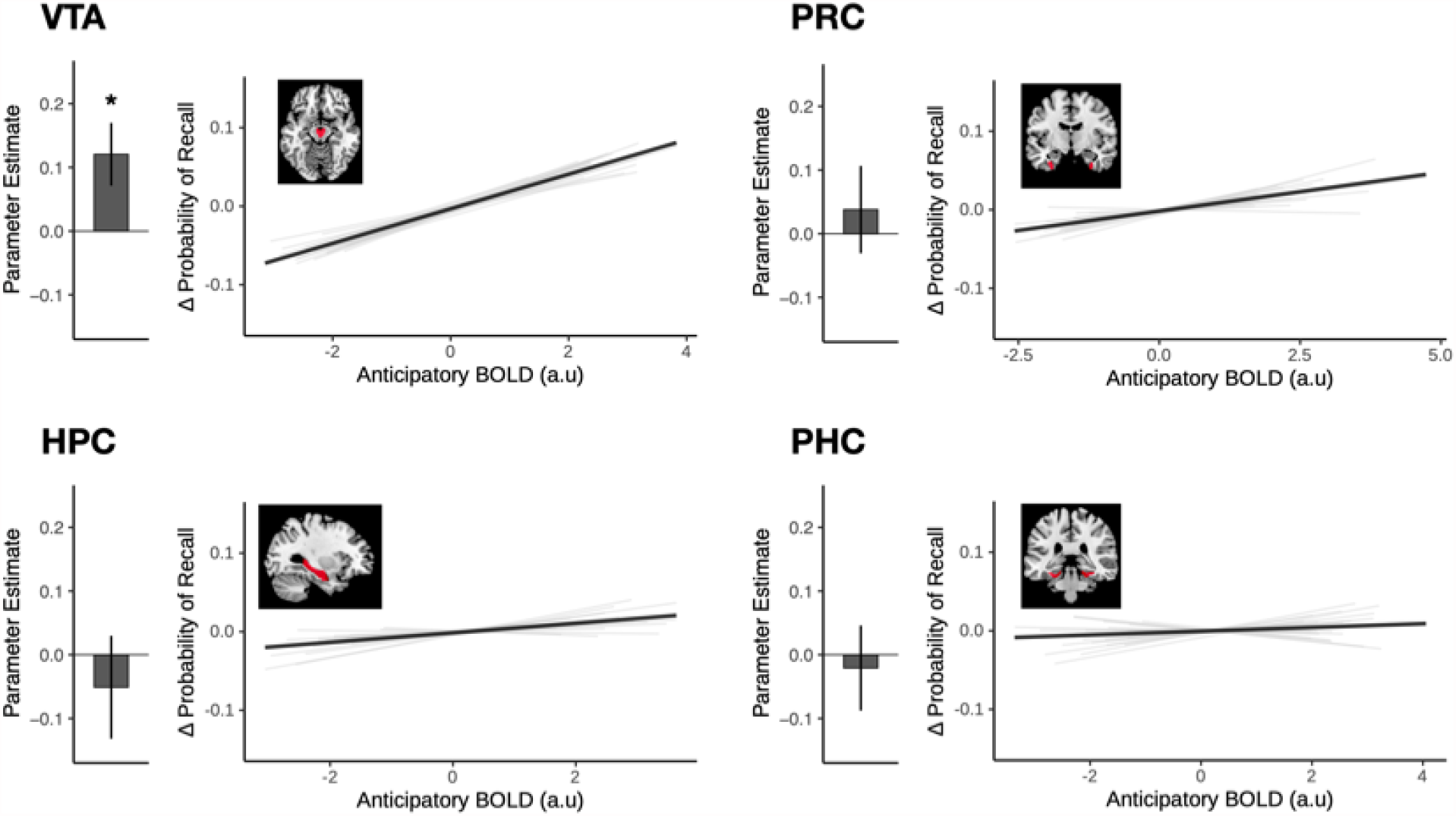
VTA univariate activation during anticipation of trivia answers uniquely explains subsequent recall. Anticipatory BOLD activation (i.e., during the Question interval preceding each trivia answer) in the VTA and medial temporal lobe ROIs was used to predict memory outcome for each trial in a mixed-effects logistic regression model. This method allows the identification of variance that is uniquely accounted for by each of the ROIs. Bar graphs in each panel represent the parameter estimate for each ROI in the full model. Among the ROIs, VTA activation was the only statistically significant predictor of subsequent recall of answers. For visualisation, the estimated predicted probability of recall relative to the individual’s mean probability (delta from within-subject mean, y-axis) is plotted against the univariate signal in each ROI (arbitrary units, a.u.; x-axis). Light gray lines depict the slope for each participant, while the solid black line depicts the mean slope across all participants. Error bars represent the SEM. ^**\***^ *p* < .05.

### Hippocampal convergence during anticipation of answers increased after high-curiosity questions and predicted better recall

Given our hypothesis that the motivation to learn would bias anticipatory hippocampal states toward a successful encoding state that followed the Anna Karenina principle, we devised an approach to characterize hippocampal states based on their pattern ‘convergence’, or the distance from an ‘average-state’ (Figure 4A; refer to *“Multivariate convergence analysis”* in the Methods section for details). For this analysis, we used a leave-one-run out approach where a cluster centroid was defined using data from N-1 runs. The multivariate activation pattern for each trial can be operationalized as a point in a high dimensional state space. The centroid is the point with the shortest distance (Pearson’s correlation distance) to all other points in the state space. This centroid was then used as the origin to quantify the distance for trials from the left-out run. This was repeated for all runs, and each trial was assigned a value representing the distance between the activation pattern for that trial, and an independently defined ‘average’ pattern. In the current formulation, patterns closer to the centroid (i.e. shorter distances), are considered to exhibit greater convergence than patterns further from the centroid.

**Figure 4.**
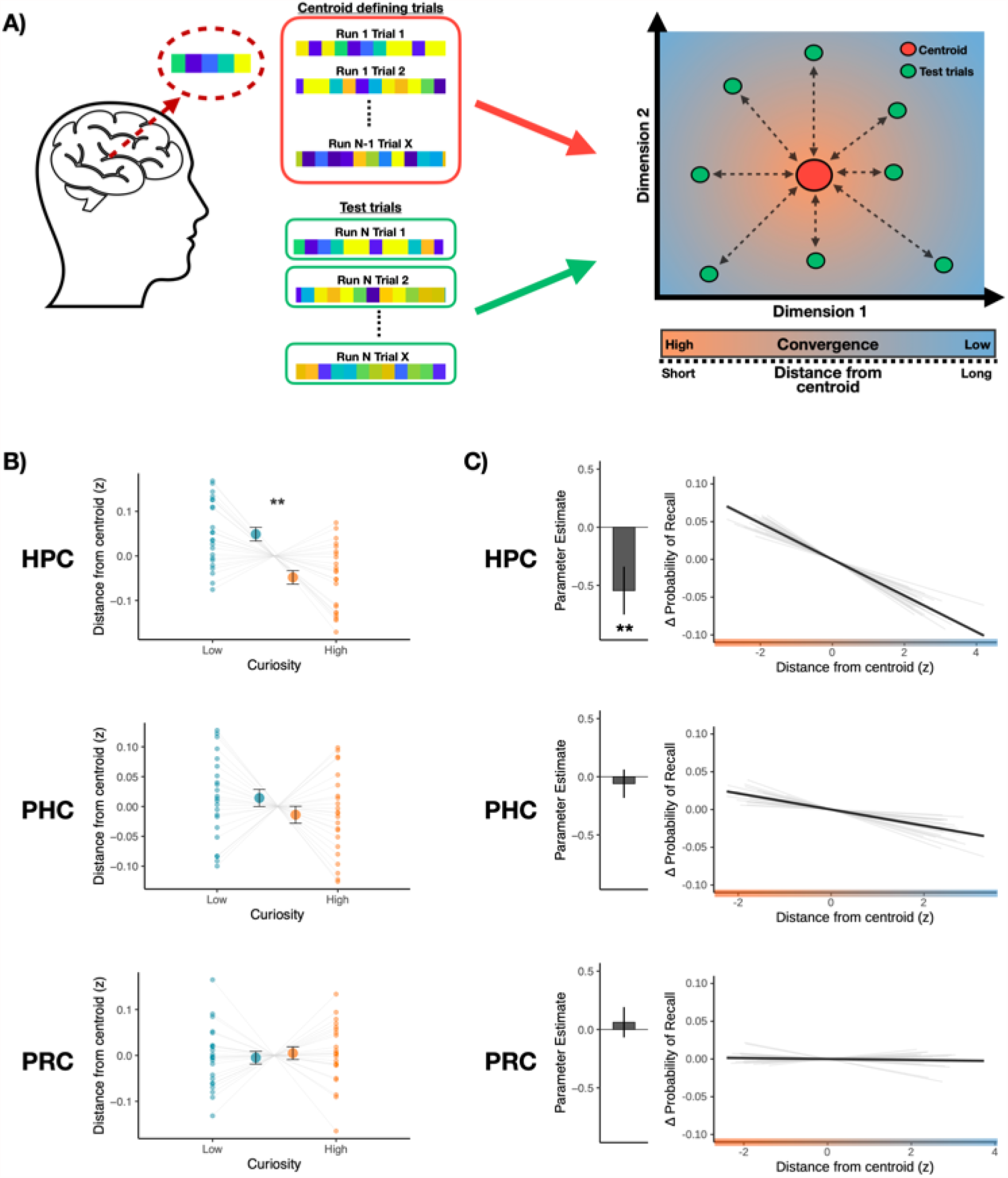
Anticipatory state convergence in the hippocampus is uniquely associated with curiosity and subsequent recall. **A)** Patterns of activation for each trial were extracted from each of the ROIs, and convergence is operationalised as the distance between the activation pattern of each trial and an independently defined centroid (i.e., an ‘average state’). A leave-one-run-out approach was used where the cluster centroid was identified using data from N-1 runs. This centroid was then used as the origin to quantify the distance for trials from the left-out run. Patterns closer to the centroid (shorter distances) are considered to exhibit greater convergence than patterns further from the centroid. **B)** During the anticipation of High-Curiosity trivia answers, patterns of activation in the hippocampus (HPC) exhibited greater state convergence, but not the surrounding medial temporal lobe cortices (PHC: Parahippocampal cortex; PRC: Perirhinal cortex). The larger dots in each panel represents the group mean, while the smaller dots represent the mean distance for each participant. **C)** We used a mixed-effects logistic regression model to predict memory outcome for each trial using the state convergence of the medial temporal lobe ROIs. Hippocampal convergence was the only significant predictor of subsequent recall. Bar graph of each panel represents the parameter estimate of each ROI in the full model. For visualisation, the estimated change in probability of recall (demeaned within subject) is plotted against the distance from centroid for each ROI. Light gray lines depict the slope for each participant, while the solid black line depicts the mean slope across all participants. Error bars represent the SEM. ^**^ *p* < .01.

We first examined whether the convergence of anticipatory states in the medial temporal lobe ROIs was influenced by curiosity (Figure 4B). Consistent with the expectation that a motivation to learn may bias hippocampal state, we observed a significant main effect of curiosity, such that spatially distributed patterns in the hippocampus showed greater convergence (i.e. shorter distance to centroid) during high-curiosity anticipation than during low-curiosity (*β* = -.017, *SE* = .007, *p* =.011). This effect was seen only in the hippocampus, not in the surrounding parahippocampal (*β* = -.009, *SE* = .011, *p* =.421) or perirhinal cortices (*β* = .003, *SE* = .010, *p* = .788).

To examine whether state convergence during anticipation predicted subsequent recall (Figure 4C), we used a mixed effect logistic regression. State convergence in the hippocampus was the only significant predictor of subsequent recall. Thus, anticipatory activation patterns that were more convergent were also associated with a higher likelihood of successful memory formation (*β* = -0.55, *SE* = .21, *p* =.008). Like the effects of curiosity, the relationship of anticipatory convergence to memory was also specific to the hippocampus, and not seen in the surrounding medial temporal cortices (Parahippocampal cortex: *β* = =.06, *p* = .622; Perirhinal cortex: *β* = .06, *p* =.634). While anticipatory univariate activation in the hippocampus was not a significant predictor of subsequent recall, we ran an additional control analysis including univariate signal in the hippocampus as a covariate to ensure that the pattern convergence effect was not driven by differences in signal amplitude. Convergence in the hippocampus remained a significant predictor after controlling for univariate activation (*β* = -0.55, *SE* = .21, *p* = .008). Moreover, the inclusion of curiosity as an interaction term improved the model fit (**χ**^2^ = 191.28, *p* < .001). Posthoc comparisons showed that the slope was significantly greater in the low curiosity condition than in the high curiosity condition (*β*_*Difference*_ = -1.24, *SE* = .09, *p* <.001), suggesting that low anticipatory hippocampal convergence may be particularly damaging to subsequent memory formation when the motivation to learn is lower.

### Trial-to-trial VTA activation predicted hippocampal convergence during anticipation of answers

Central to our primary hypothesis, univariate activation in the VTA during anticipation of answers was a significant predictor of convergence in the hippocampus (*β* = -.059, *SE* = .012, *p* < .001, Figure 5A), with greater VTA activation associated with greater hippocampal convergence (i.e. shorter distances). This result remained robust when univariate hippocampal activity was included as a covariate (*β* = -.049, *SE* = .011, *p* < .001). To ensure that this was not simply driven by differences between the curiosity states, a model comparison was performed comparing model fit between models with and without an interaction term for the Curiosity condition. The inclusion of an interaction term did not result in a better model fit (**χ**^2^ = 0.44, *p* = .507), suggesting that the association between VTA activation and hippocampal convergence was not driven by differences across the curiosity states.

**Figure 5.**
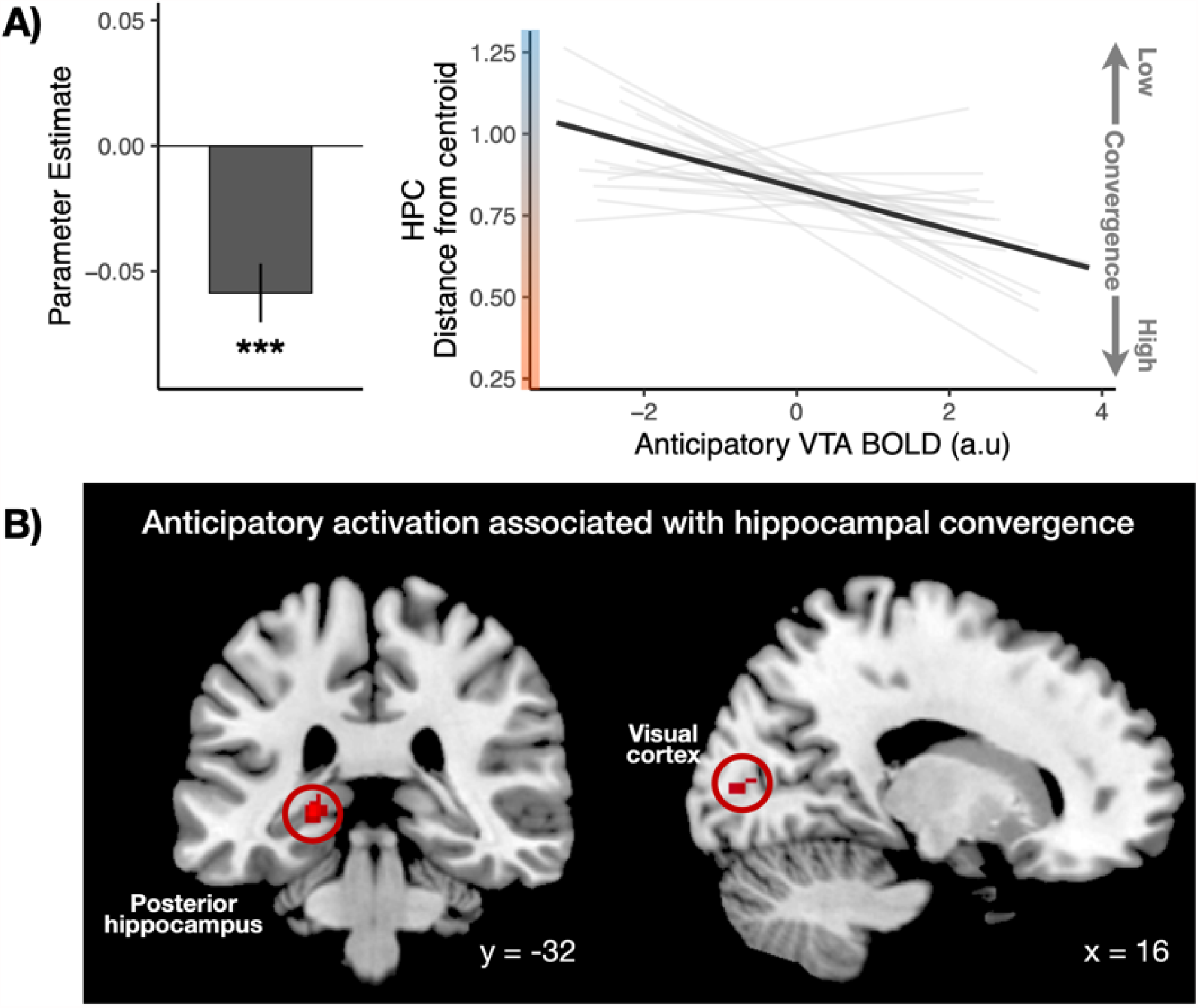
Anticipatory VTA activation selectively modulates hippocampal state convergence. **(A)** Greater activation in the VTA is associated with greater convergence in the hippocampus during the anticipation of trivia answers. The bar graph represents the parameter estimate for the association of VTA activation and hippocampal distance. For visualisation, the predicted distance from centroid for the hippocampus is plotted against univariate VTA BOLD activation. Light gray lines depict the slope for each participant, while the solid black line depicts the mean slope across all participants. Convergence is defined as a shorter distance from centroid. **(B)** Whole-brain analysis (controlling for univariate VTA activation) showed that anticipatory univariate activations in clusters that included the visual cortex and posterior hippocampus are also positively associated with convergence in the HPC. Error bars represent the SEM. ^***^ p< .001.

To examine whether convergence in the hippocampus was also associated with univariate activation in other brain regions, we performed an exploratory whole-brain voxel-wise analysis correlating hippocampal convergence with each voxel’s univariate activation (controlling for VTA activation). To control for spurious correlations, a null distribution was generated for each voxel using a permutation-based approach (500 iterations), and the r-value at the 95th percentile was subtracted from each voxel. This approach ensured that only correlation values greater than the 95th percentile of the null distribution are positive, and a one-sample t-test was then performed across subjects to identify regions showing a significant correlation with hippocampal convergence. At a statistical threshold of *p* < .05 (FWE-corrected), we see significant clusters only in the posterior hippocampus (Peak coord: -22, -32, -2, t = 10.90; Figure 5B) and the early visual cortex (Peak Coord: 16, -86, 6, t = 7.49).

### Hippocampal convergence uniquely accounted for the effect of anticipatory VTA activation on memory recall

Based on our prediction that activity in the VTA influences subsequent learning through its influence on hippocampal state, we compared mixed-effects logistic regression models that included either VTA activation, hippocampal convergence, or both terms. Consistent with our hypothesis, the inclusion of both VTA activation and hippocampal convergence resulted in a better fit than a model with only VTA activation (**χ**^2^ = 5.19, *p* = .023), but not a model with only hippocampal convergence (**χ**^2^ = 2.81, *p* = .09). This was further supported by a mediation analysis showing a significant mediation of VTA activation by hippocampal convergence (pME = .276, 95% CI = [.028 1.09], *p* = .032). Together, these findings suggest that the influence of anticipatory VTA activation on subsequent recall is primarily mediated via specific effects on neural state in the hippocampus.

### Individual differences in VTA activation and Hippocampal convergence

Complementing the intra-individual analyses reported, we also observed a significant correlation of VTA activation and hippocampal convergence across individuals, where participants with greater VTA activation showed greater convergence in the HPC (r = -.56, 95% CI = [-.79 .19], *p* =.005). This relationship was observed for both the High curiosity (r = -.57, 95% CI = [-.72 -.22], *p* =.004) and Low curiosity conditions (r = -.44, 95% CI = [-.72 -.04], *p* = .033), suggesting that beyond intra-individual variation, individual difference in the engagement of VTA is also associated with hippocampal convergence.

Additionally, we examined if individual difference in memory performance is associated with curiosity-related modulation of hippocampal convergence. While we did not observe a significant correlation between the difference in curiosity-related hippocampal convergence (Low - High curiosity) and memory benefits (r = -.05, 95% CI = [-.45 .37], *p* = .831), we did observe a significant correlation between curiosity-related hippocampal convergence and overall memory performance (across both conditions) whereby a greater curiosity-related hippocampal convergence was associated with better memory performance (r = .44, 95% CI = [.03 .72], *p* = .037).

### Encoding-related convergence in the medial temporal cortices during answers predicted subsequent recall unrelated to curiosity

While our primary focus surrounds the anticipatory state following the presentation of the trivia questions, for completeness, we conducted the same analyses for activation associated with the presentation of trivia answers. High curiosity was associated with a trend towards greater activation in the VTA (*β* = .059, *SE* = .031, *p* = .057). None of the medial temporal lobe ROIs showed a significant effect of curiosity during the presentation of answers (Hippocampus: *β* = -.020, *SE* = .028, *p* = .481; Parahippocampal cortex: *β* = -.017, *SE* = .029, *p* = .574; Perirhinal cortex: *β* = .036, *SE* = .024, *p* = .127). Across all ROIs, only activation in the perirhinal cortex was associated with a greater likelihood of recall (*β* = .223, *SE* = .076, *p* = .003). These findings held when activation across both intervals (Question & Answers) were included in a single model (Supplementary Figure 1), suggesting that univariate activation during the presentation of questions and answers account for unique variance related to memory outcomes.

We also performed analyses of pattern convergence during the presentation and encoding of trivia answers (Supplementary Figure 2). Curiosity did not predict state convergence during answers in any of the medial temporal lobe ROIs (Hippocampus: *β* = .002, *SE* = .008, *p* =.750 ; Parahippocampal cortex: *β* =.006, *SE* = .011, *p* =.558; Perirhinal cortex: *β* = -.009, *SE* = .010, *p* = .361). Convergence during answers in the parahippocampal (*β* = -.500, *SE* = .133, *p* < .001) and perirhinal cortices (*β* = -.605, *SE* = .139, *p* <.001) but not the hippocampus (*β* =.124, *SE* = .184, *p* = .500), significantly predicted subsequent recall, unrelated to curiosity. This remained significant after controlling for univariate activation in both the parahippocampal (*β* = -.379, *SE* =.132, *p* = .004) and perirhinal cortices (*β* = -.558, *SE* = .137, *p* < .001).

## Discussion

The present findings identify midbrain VTA neuromodulation of anticipatory hippocampal state convergence as a candidate mechanism of motivated memory. By using multivoxel pattern analysis on fMRI data acquired during anticipation of answers associated with high or low curiosity, we related univariate VTA activation to states in the medial temporal lobe. After trivia questions eliciting high curiosity, patterns of activation in the hippocampus, but not the surrounding medial temporal cortices, were biased towards greater convergence (i.e. lower variability), and anticipatory hippocampal convergence was uniquely associated with later recall of the anticipated answers. Across individuals, curiosity-related modulation of hippocampal convergence was correlated with recall. Most importantly, convergence in the hippocampus was strongly associated with trial-by-trial variation in anticipatory VTA activation and uniquely accounted for the relationship between VTA activation and subsequent recall. We conclude that higher curiosity during anticipation of answers engaged VTA activation, stabilized anticipatory convergence states specifically in the hippocampus, and thus enhanced memory formation. This observed cascade offers potential answers to longstanding questions about neuromodulation of memory and about fundamental hippocampal mechanisms.

Our approach leverages prior work in which fMRI multivoxel pattern analysis was used to examine representations of content during encoding. In prior studies, similarity in neural patterns across repeated occurrences of the same stimulus was associated with better memory, attributed to reinstatement of stimulus specific information (Poh and Chee, 2017; Xue et al., 2010, 2013). Similar analytical approaches comparing patterns of activation across consecutive timepoints have also been used to examine temporal dynamics in the hippocampus (Brunec et al., 2018). Here, we used pattern analysis not to study stimulus representations or temporal dynamics, but rather to identify consistent engagement of states or cognitive processes associated with the anticipation of answers and successful memory formation. Our measure, convergence, is defined based on the distance from an average activation pattern that includes trials from all conditions, so our finding implies that patterns associated with subsequent remembering exhibit greater ‘typicality’, while patterns associated with forgetting show greater variability. This dissociation is consistent with the expectation that successful memory formation is likely to require the convergence of multiple cognitive operations and physiological conditions, while memory failures can arise from disruption to any of the component elements, following the Anna Karenina principle. In the domain of human neuroimaging, this principle has been applied in large-scale network analysis to examine individual differences (Finn et al., 2020). Here, by showing that hippocampal convergence was positively associated with both activation in the midbrain and with subsequent recall (Figure 6), we demonstrate that the Anna Karenina principle can similarly be applied to examine intra-individual variability in cognitive states to predict momentary behavior.

**Figure 6.**
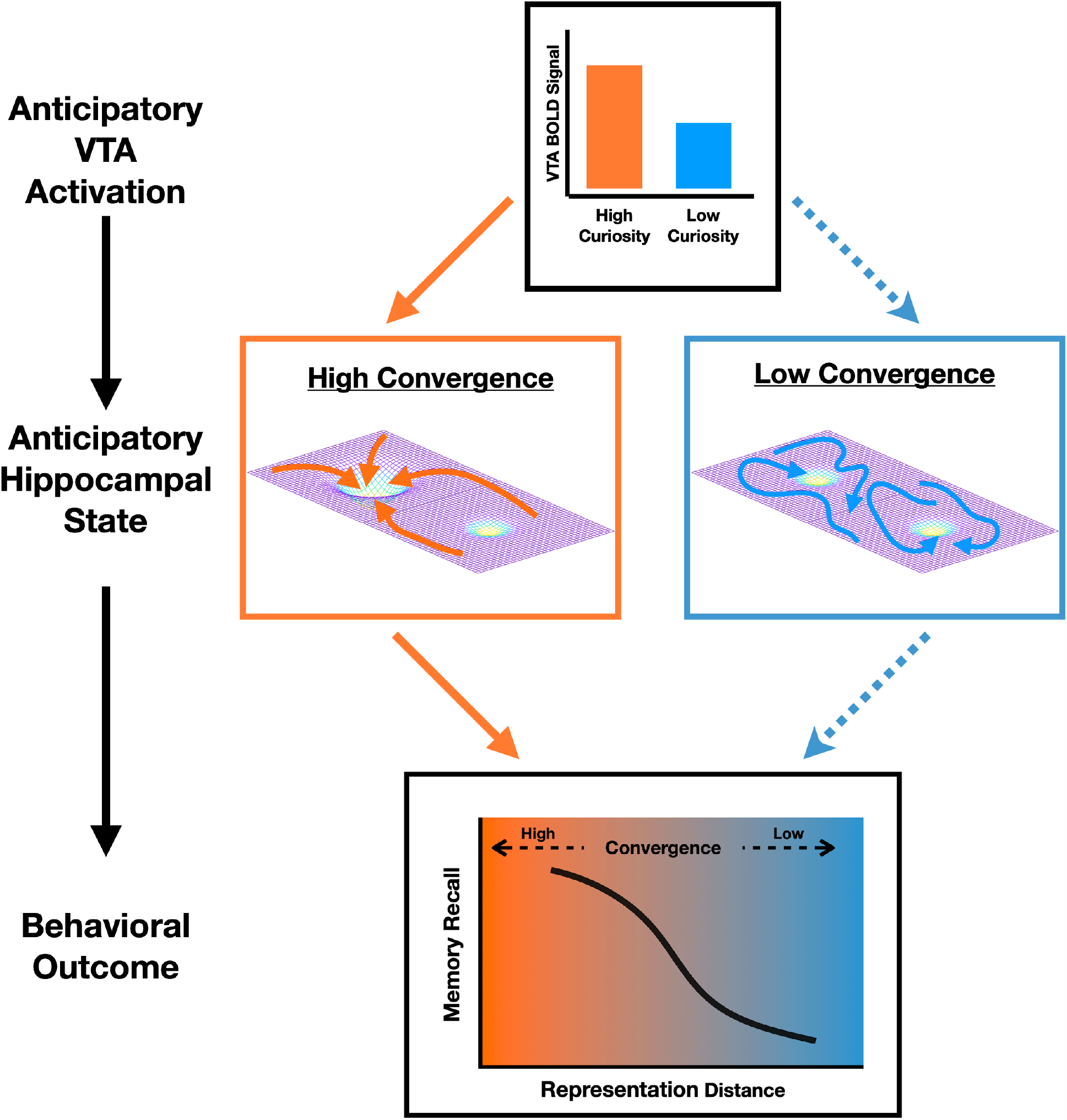
Schematic depiction of Hippocampal convergence with neuromodulation from the VTA. During the anticipation of high curiosity answers, neuromodulation by the VTA promotes the consistent engagement of neural states/processes in the hippocampus that supports the formation of new memories. This can manifest as greater convergence in distributed patterns of activity. In the absence of neuromodulatory inputs from the VTA (during low curiosity), patterns of activity in the hippocampus can show greater variability. This increased convergence in hippocampal state with VTA neuromodulation may be supported by the stabilization of specific attractors and the suppression of noise-driven transitions between different possible attractor states.

While our current approach is agnostic to underlying processes manifesting in a convergence state, a central question that remains is how an anticipatory convergence state in the hippocampus specifically benefits the formation of new memories. We consider two potential mechanisms. First, it has been suggested that the hippocampus can maintain multiple spatial maps of the environment, allowing for dynamic switching between spatial frames depending on the relevant context (Kelemen and Fenton, 2016). Here, a ‘contextual frame’ account would suggest that convergence in the hippocampus may reflect the instantiation of a ‘motivated context /map’ that serves as the scaffold in which new memories may be embedded. This instantiation of a specific reference frame may be akin to processes underlying the ‘Memory palace’ or the method of loci mnemonic device, where memorization is supported through the active instantiation and maintenance of a spatial reference frame that accommodates the ‘storage’ of to-be-remembered information. An alternative from dynamic systems is an optimal subspace account. In studies of voluntary movement, it has been suggested that preparatory activity in the motor cortex may reflect the initialization of a neural state within an optimal subspace, while an evolving neural trajectory encodes the motor execution (Churchland et al., 2006, 2010a). In the memory domain, information to be encoded may be similarly represented as a neural trajectory that evolves from an anticipatory state, with trajectories originating from an optimal subspace predicting memory formation. High trial-to-trial variability in initial state could increase overlap of the subsequent encoding trajectories, reducing the distinctiveness of memory traces (Zylberberg et al., 2016). Much in the same way that hikers may halt a conversation when attempting to locate the sound of a flowing stream, a convergent state may thus reduce noise-driven variability, supporting more efficient encoding of the incoming signal.

Related lines of work on spontaneous brain dynamics have consistently shown that variability in neural response can be reduced by exogenous stimulus (Churchland et al., 2010b) or by endogenous allocation of attention (Arazi et al., 2019; Cohen and Maunsell, 2009). This reduction in neural variability has been associated with enhanced processing of stimulus-related information (Schurger et al., 2015), and greater sensitivity in perceptual performance (Arazi et al., 2019). Computational accounts have proposed that the underlying mechanism of such reduction in neural variability may be the stabilization of specific attractors and the suppression of noise-driven transitions between different possible attractor states (Deco and Hugues, 2012). Though it may be tempting to attribute our current observation to similar underlying mechanisms, there are several key differences that should be noted. While the reduction in neural variability has been observed across widespread brain regions at different levels of analysis (e.g. extracellular recording (Churchland et al., 2010b), MRI (He, 2013), ECoG (He and Zempel, 2013), EEG (Arazi et al., 2017)), these are primarily observed in neocortical regions and it is uncertain if the same phenomenon would manifest in the hippocampus. In our current study, hippocampal convergence was modulated by curiosity and was predictive of subsequent recall only during anticipation (Question interval), whereas convergence in the medial temporal cortices was predictive of memory only during the presentation of information (Answer interval). These findings are consistent with an account of enhanced information processing with reduced neural variability, but also raises an intriguing possibility that convergence in the hippocampus may be more strongly modulated by internal states (curiosity in this case), whereas convergence in the medial temporal cortices may be more strongly associated with attention to external stimulus.

The idea that internal motivational states like curiosity modulate hippocampal function has now been substantiated by accumulating evidence that midbrain dopaminergic circuitry interacts with the hippocampus across multiple timescales to influence the formation of episodic memory. Neurobiological mechanisms that have been proposed to explain the effects of dopamine on memory formation in the hippocampus include meta-plastic changes, such as lowering the threshold for LTP (Li et al., 2003; Otmakhova and Lisman, 1996), or ‘tagging’ for subsequent consolidation (Redondo and Morris, 2011). It is unclear how, or whether, such meta-plastic changes would translate to metabolic demand, and thus changes in univariate BOLD activity during encoding. However, neuromodulation by dopamine can also alter the physiological properties of the hippocampus. We reasoned that while an fMRI analytical approach cannot provide evidence about meta-plastic changes not reflected in BOLD, it is well-suited to detect modulatory influences that establish a distributed state across the hippocampus conducive to memory formation, including the convergence state we hypothesized.

It is known that the hippocampus can exhibit distinct functional states that influence how information is processed and subsequently encoded (Carr and Frank, 2012; de Chastelaine and Rugg, 2015; Fell et al., 2011; Guderian et al., 2009; Kennedy and Shapiro, 2009; Richter et al., 2016; Urgolites et al., 2020; Wolosin et al., 2013; Zeithamova et al., 2018). Computational models of episodic memory have proposed that the hippocampal memory system may alternate between functional states supporting encoding or retrieval (Decker and Duncan, 2020; Hasselmo et al., 1996, 2002; Meeter et al., 2004; Paulsen and Moser, 1998). While alternation between encoding and retrieval states has been suggested to occur on a rapid timescale based on theta phase, evidence from both human (Kirov et al., 2009; Long and Kuhl, 2019) and rodent (Molter et al., 2012) studies have also shown oscillatory signatures of sustained states aligned with the timescales of neuromodulatory influence. Such sustained state changes have similarly been observed using fMRI, where patterns of activation were successfully used to classify states associated with encoding or retrieval (Richter et al., 2016), and in identifying fluctuation of network states relating to encoding success (Keerativittayayut et al., 2018). In addition, behavioral studies have also shown evidence that hippocampal processing can fluctuate between a bias towards pattern separation or completion over several seconds (Duncan et al., 2012; Patil and Duncan, 2017). As our current analytical approach focuses on anticipatory convergence states, it is unclear if our observation corresponds to any of the previously identified states, or to a theoretical ‘encoding-state’. However, it should be noted that the current task requires not only encoding of the trivia answer, but also binding of the answer to the associated question. Although we have no separate measures of item encoding and relational binding, we saw a dissociation consistent with their distinct roles (Davachi, 2006; Davachi and Mitchell, 2003; Davachi and Wagner, 2002) in item and relational memory: recall was predicted by hippocampal convergence bridging the gap between the question and answer, but by perirhinal convergence during presentation of answers alone. The observation of different intervals for involvement of hippocampal and perirhinal convergence also speaks against a purely attentional account of our findings, as the maintenance of attention should be sustained across time, manifesting in both intervals.

In human fMRI studies, overall increases in activation of the midbrain and the hippocampus preceding (Adcock et al., 2006; Gruber et al., 2014; Wittmann et al., 2005), during (Aberg et al., 2020; Loh et al., 2016; Ripollés et al., 2016; Wolosin et al., 2012), and following encoding (Gruber et al., 2016; Murty et al., 2017; Tompary et al., 2015) have been related to better subsequent memory performance. While these prior findings are consistent with an account of dopaminergic neuromodulation, the mechanisms of such effects have remained to be specified. As discussed above, some known mechanisms of dopamine on hippocampal plasticity (metaplastic tag and capture, lowered LTP threshold) might not manifest in univariate BOLD activation. In particular, it was previously unknown how or whether spatially distributed patterns of activity in the hippocampus are modulated by midbrain activity. Our work bridges these gaps, further substantiating an account of neuromodulation of the hippocampus by midbrain activity, by showing a trial-level association between activation in the midbrain and convergence of distributed patterns in the hippocampus.

In contrast to the results of prior studies showing greater univariate activation in the hippocampus preceding successful memory formation (Adcock et al., 2006; Gruber et al., 2014), we observed a relationship between subsequent memory not with univariate signal magnitude, but with multivariate pattern convergence. Apart from key differences in experimental design that could have influenced anticipatory activity in prior studies, such as the use of intentional encoding (Adcock et al., 2006), or the expectation of irrelevant faces (Gruber et al., 2014), several analytical decisions may also have contributed to the difference in observations. In particular, our current univariate analysis is performed on the average signal across all voxels within our ROI, and was not optimized to isolate localised univariate differences. Additionally, in both of the prior studies, the mnemonic effect in the hippocampus was specific to the high motivation condition, whereas here, we showed that multivariate convergence in the hippocampus was associated with subsequent recall in both the high and low curiosity states, suggesting that distributed patterns of activity may be more sensitive for the detection of such mnemonic effects. However, it should be noted that multivariate convergence and univariate event-related activation can co-occur, as evidenced in the medial temporal cortices during answer presentation, where both univariate activation and multivariate convergence were associated with subsequent recall. Furthermore, convergence in medial temporal cortices remained a significant predictor of recall when controlling for univariate activity, suggesting that these measures have dissociable mechanisms contributing to memory formation. We did not observe an association between anticipatory midbrain and anticipatory hippocampal univariate activation. Prior findings aside, this should not be entirely surprising, because not all neuromodulatory effects that promote memory formation should be evident in changes in magnitude of BOLD activation; this premise is a main motivation for the current study. An optimal subspace account could explain prior findings of increased anticipatory univariate activation within the hippocampus: Such activation would be expected under conditions where greater metabolic activity is required to ‘shift’ ongoing states into the optimal subspace.

Our current contribution focuses on evidence of neuromodulation of the hippocampus by dopaminergic nuclei in the midbrain. It should be noted that activation of the midbrain is also associated with the release of other neurotransmitters besides dopamine, and the hippocampus is also innervated by other major neuromodulatory nuclei including the basal forebrain, the primary source of cholinergic projections. The release of acetylcholine in the hippocampus has been shown to promote memory formation, and it has been suggested that the fluctuations in levels of acetylcholine can drive functional states in the hippocampus, with high acetylcholine promoting an encoding state and pattern separation, while low acetylcholine may promote a retrieval state and pattern completion (Decker and Duncan, 2020; Hasselmo, 1999; Hasselmo et al., 1995). Although our exploratory whole-brain analysis did not reveal a significant association in the basal forebrain, this could be due to a lack of sensitivity in our current approach. It should be noted that the noradrenergic locus coeruleus (LC) has also been shown to release dopamine in the hippocampus (Kempadoo et al., 2016; Takeuchi et al., 2016), and recent work in mice has suggested that novelty-enhanced memory may be more strongly dependent on projections from the LC than the VTA (Takeuchi et al., 2016). While our current acquisition precludes a precise localization of the LC, future work should also examine if activity in the LC is similarly associated with hippocampal convergence. If hippocampal convergence is an outcome of dopaminergic modulation irrespective of its source or dynamics, then many manipulations, including novelty and prediction error, should all promote greater hippocampal convergence. We have argued previously that mesolimbic signalling to the hippocampus is more likely to be sustained than phasic (Shohamy and Adcock, 2010; Murty et al., 2016). An intriguing possibility is that hippocampal convergence is a memory mechanism specific to modulation by dopamine from mesolimbic projections.

Finally, while we interpret the VTA-Hippocampal interaction here from a neuromodulatory perspective, with hippocampal convergence as an outcome of VTA modulation, it should be noted that activity in the VTA is also influenced, indirectly, by signalling from the hippocampus (Floresco et al., 2001; Lisman and Grace, 2005). We have previously modeled VTA activation in fMRI data using this reciprocal relationship (Murty et al., 2016). Further studies will be necessary to clarify the directionality and neurotransmitter specificity of these anatomical relationships. A main contribution of the present work is a candidate physiological manifestation of neuromodulation, to be tested via further investigation in both human and animal models.

## Conclusion

Using a novel systems-level characterization of hippocampal function, a stable convergence state, we demonstrate a potential mechanism of mesolimbic neuromodulation that can account for its effects on memory formation. The present study suggests that activation of the midbrain VTA by curiosity promotes an anticipatory convergence state specific to the hippocampus that may be optimal for the encoding of new information. These findings set the stage for future work to understand distinct neural signatures of functional states and neuromodulatory system effects. Broadly speaking, the concept of a convergence state could hold promise to not only illuminate modulation of memory, but also potentially unite piecemeal understanding of individual mechanisms into a cohesive picture of the role of the hippocampus in memory formation.

## Methods

### Subjects

Twenty-five healthy, right-handed young adults were recruited for the study. All participants provided informed consent for our study protocol approved by the Duke University Institutional Review Board. Two participants had to be excluded (one participant fell asleep during the scan, and one did not complete the scanning session), and all remaining 23 participants were included in the analyses (10 female; Mean age = 26.4 years, Age range = 19-35 years).

### Tasks

#### Trivia question stimulus screening

We selected 360 trivia questions from the stimuli used in the study by Gruber and colleagues (2014), and a pre-task screening session was used to sort trivia questions into high- and low-curiosity categories for each participant. For the pre-screening session, participants were presented with a series of trivia questions and they were required to make self-paced ratings on a continuous scale. Participants responded to the following questions: 1) “How likely is it that you know the answerã” and 2) “How curious are you about the answerã”. Trivia questions were excluded if participants indicated a high likelihood of knowing the answer (>90% on the scale), and they responded until 216 trivia questions were eligible for inclusion. Included trivia questions were separated into tertiles based on curiosity ratings (72 questions each), with questions in the 1st and 3rd tertiles categorized as low and high curiosity respectively. Twelve questions from the 2nd tertile were used as catch trials during encoding and were not included in any analysis.

#### Incidental encoding of trivia questions and answers

Participants performed the encoding task during fMRI scanning where they were shown the trivia questions (Question Interval) and were presented with the associated answer (Answer Interval) after a variable delay interval. During the question presentation, participants were shown a single trivia question together with a colored rectangle for 4 seconds. The colored rectangle indicated the duration and action contingency for the upcoming trial, whereby the length of the rectangle indicated the duration of the anticipation period (9s or 13s), and the color indicated if participants were required to make a button press to see the trivia answer. On trials that required a button press, a green arrow appeared on the left or right side of the screen, and participants made a button press to indicate the side that the arrow was presented on. For action contingent trials, the trivia answer was shown only if the participants responded correctly or the string ‘XXXXX’ would be presented (Participants were highly accurate and saw the trivia answer for most trials (M = 98.6%, SD = 0.4). Only trials that were correctly responded to were included for subsequent analyses). On non-action contingent trials, the trivia answer was shown at the end of the anticipation period. The manipulation of action-contingency was performed for the investigation of a separate research question that would be elaborated on in a separate communication. The trivia answer was presented for 1 second, following which, participants performed an active baseline task where they were required to count backward from different starting numbers for a duration between 1 to 20 seconds. To encourage compliance with the active baseline task, catch trials occurred at random intervals, and participants were required to indicate whether their current count was above or below a given number. There were a total of 12 catch trials and trivia questions presented following the catch trials (taken from the 2nd tertile) were not included in the analysis.

Participants underwent a total of 6 scanning runs (10 mins each), with 12 high curiosity trials, 12 low curiosity trials, and 2 catch trials presented within each run. Within each curiosity condition, there was an equal number of action contingent and non-action contingent trials. Condition onset and trial intervals were optimized using OptSeq2 (Dale, 1999).

#### Surprise Recall

Immediately following the scan, participants were given a surprise recall test for the trivia questions. Participants were shown all 144 (72 High and 72 Low curiosity) trivia questions in random order, and were required to type out the correct answer for each question. Participants were told not to make any guesses if they were unable to remember the correct answer.

### MRI Data Acquisition

MRI data were acquired on a 3T GE Signa MRI scanner at the Duke Brain Imaging and Analysis Center. fMRI data for each participant were acquired using an echo-planar imaging (EPI) sequence (TE = 27ms, flip angle = 77 degrees, TR = 2000ms, voxel size = 3.75mm x 3.75mm) with 34 axial slices (slice thickness = 3.8mm). Participants completed a total of 6 functional scan runs each consisting of 298 fMRI volumes. Cardiac and respiratory physiological data were also collected during functional scans using BioPac. Prior to the functional scans, whole-brain, inversion recovery, spoiled gradient high resolution anatomical image (voxel size = 1mm isotropic) was collected for spatial normalization.

#### fMRI Preprocessing

Preprocessing of the fMRI data was performed using fMRI Expert Analysis Tool (FEAT) Version 6.00 implemented on FSL 5.0.8 (www.fmrib.ox.ac.uk/fsl). The first 6 volumes from each scan run were discarded to allow for signal stabilization. Physiological noise correction was performed using the Physiological Noise Modeling toolbox in FSL. Skull stripping was performed using BET (Jenkinson et al., 2005), and images were realigned within-run, intensity normalized by a single multiplicative factor, spatially smoothed with a 4 mm full-width half-maximum (FWHM) kernel, and subjected to a high-pass filter (80s). The 4mm smoothing kernel was chosen to optimize the differentiation of midbrain and hippocampal signals (Adcock et al., 2006). Spatial normalization was performed using a two-step procedure using FLIRT, where mean EPI from each run was co-registered to the high-resolution anatomical image, which was followed by the normalization of the high-resolution anatomical image to MNI space using a nonlinear transformation with a 10mm warp resolution.

#### Defining regions-of-interest

To examine how activity in the midbrain interacts with the medial temporal lobe (MTL), we identified regions of interest which included the midbrain VTA and regions within the MTL. The VTA was defined using a probabilistic atlas thresholded at 50% (Murty et al., 2014). Three separate ROIs were defined within the medial temporal lobe, which included the hippocampus proper, perirhinal cortex and parahippocampal cortex. The hippocampus was defined using the Harvard-Oxford structural atlas, while the perirhinal and parahippocampal cortex were defined using anatomical mask from (Ritchey et al., 2015). All ROIs were defined in MNI space.

### Analysis

Due to a programming error, trivia questions for one participant were not correctly selected (based on screening), and 62 trials were removed from subsequent analysis. Statistical analysis was performed using linear and logistic mixed-effects modeling using the lme4 (Bates et al., 2015) and lmerTest (Kuznetsova et al., 2017) packages in R (R Core Team, 2020). Data visualization was generated using the ggplot2 package (Wickham, 2016).

#### Behavioral Analysis of Memory performance

Memory performance was analyzed using a paired t-test comparing recall rates between the high and low curiosity condition. An additional analysis was also conducted using repeated measure ANOVA with curiosity and action-contingency as factors. Post-hoc comparisons were conducted based on the contrast of estimated marginal means, and recall rate was greater for the high curiosity condition across both levels of action-contingency.

#### fMRI Analysis

To capture trial-level estimates, we used the Least-squares separate approach (Mumford et al., 2012) to estimate the betas associated with each condition at each trial. Briefly, each trial is estimated using a separate model where the trial of interest is modeled as a separate regressor from all other trials. Separate parameter estimates were obtained for the question interval and during the answer interval, resulting in a total of 288 separate models (144 trials x 2 intervals). Parameter estimates were converted to t-values and normalized to z-values. Voxel values from the z-maps were used for both univariate and multivariate analyses.

#### Univariate analysis - Effects of Curiosity on anticipatory activity

Univariate analyses were conducted using the mean value across all voxels within each ROI during the anticipation of trivia answers (following Question presentation). Linear-mixed effects analysis was conducted for each ROI with curiosity state as fixed effect, and subjects as random effect. As previously mentioned, the manipulation of action-contingency was performed for the investigation of a separate research question, and for the current study, action-contingency was omitted from all models to increase statistical power given the limited number of trials. However, it should be noted that there was neither a main effect nor interaction of action-contingency in any of our ROIs, and the inclusion of action-contingency as a covariate did not alter any of our findings. Post-hoc comparisons were performed on the estimated marginal means using the *emmeans* package.

#### Multivariate convergence analysis

For the proposed analysis, patterns of activation in the ROIs are operationalised as points in an N-dimensional space, with N being the number of voxels in each ROI. Distance in the current analysis was measured using correlation distance (1 - Pearson’s r), a distance metric commonly used in multivoxel pattern analysis. To examine the association between VTA activity and neural state in the medial temporal lobe, we devised an approach to quantify the trial-by-trial variation in neural state based on their distance from an independently defined centroid. The cluster centroid is a point with the shortest distance to all other points in high dimensional state space, and can be thought of as an ‘average-state’. We defined the centroid using a leave-one-run-out approach, where the cluster centroid was identified, with a k-means algorithm, using data from N-1 runs. This centroid was then used as the origin to quantify the distance for trials from the left-out run. Centroids for the analysis of the anticipatory period were defined using activation patterns from the Question interval, while analysis for the encoding of answers were defined using activation patterns from the Answer interval. This was repeated for all runs and was performed independently for each subject. The convergence for each trial was quantified based on their distance from the independently defined centroid. As the trials being measured do not contribute to the definition of the centroid (which they are measured relative to), this approach ensures the independence of the tested trials and the centroid-defining samples. Additionally, this also ensures that the quantification of convergence is not confounded by temporal correlation (since the centroid is defined using data from a different scanning run). In the current formulation, patterns closer to the centroid (i.e. shorter distance) are considered to exhibit greater convergence than patterns further from the centroid. This operationalization is similar to measures of neural variability, whereby a larger absolute difference from the average signal amplitude is considered to reflect greater trial-to-trial variability (e.g. He and Zempel, 2013). In contrast to a conventional linear classification approach, which would be suboptimal given the small number of datapoints and the imbalance between conditions in the current study (between number of Remembered and Forgotten trials), this approach also capitalizes on the expectation that successful memory formation is likely to require the consistent convergence of multiple factors, while the lack of any could impede memory formation.

#### Relating univariate activity and multivariate convergence

To examine if multivariate convergence in the hippocampus is associated with univariate activity in the midbrain VTA, a linear mixed effects model was implemented with trial-level univariate activation as a predictor of hippocampal convergence. The model included subjects as random intercepts, and VTA activity as a random slope. Mediation analysis was performed using the *mediation* package.

To examine if convergence in the hippocampus is also associated with univariate activation in other brain regions, we performed an exploratory whole-brain voxel-wise analysis correlating hippocampal convergence with each voxel’s univariate activation (controlling for VTA activation). To control for spurious correlations, a null distribution was generated for each voxel using a permutation-based approach (500 iterations), and the r-value at the 95th percentile was subtracted from each voxel. This approach ensured that only correlation values greater than the 95th percentile of the null distribution are positive. A one-sample t-test was implemented using SPM12 (https://www.fil.ion.ucl.ac.uk/spm/), to identify regions showing a significant correlation with HPC convergence across all subjects. Significant voxels were identified using a threshold of FWE *p* <.05.

#### Relating brain activity and memory outcomes

To examine the behavioral relevance of univariate and multivariate measures of brain activity, mixed effects logistic regression was performed with trial-level brain measures (i.e. univariate activation or multivariate state convergence) of all ROIs included as predictors of subsequent recall. By including all ROIs in a single model, this approach allows the identification of variance that is uniquely accounted for by the activity of each ROIs. For all mixed-effects models, subjects were included as random intercepts, and random slopes were included if it generated a better model fit based on model comparisons evaluated using a likelihood ratio test.

## Data and code availability

Matlab code used for convergence analyses will be made available at - https://github.com/JiaHou-Poh/ConvergenceState. The dataset for the current study is available upon reasonable request.

## Acknowledgments

We thank Allie Sinclair, Dr. Vishnu Murty, and Dr. Denise Cai for helpful comments on the manuscript.

This work was supported by R01 MH094743 (RAA), R01 MH087610 (TE), and the NUS Development grant (JHP).

## Supplementary Materials

**Supplementary Figure 1.**
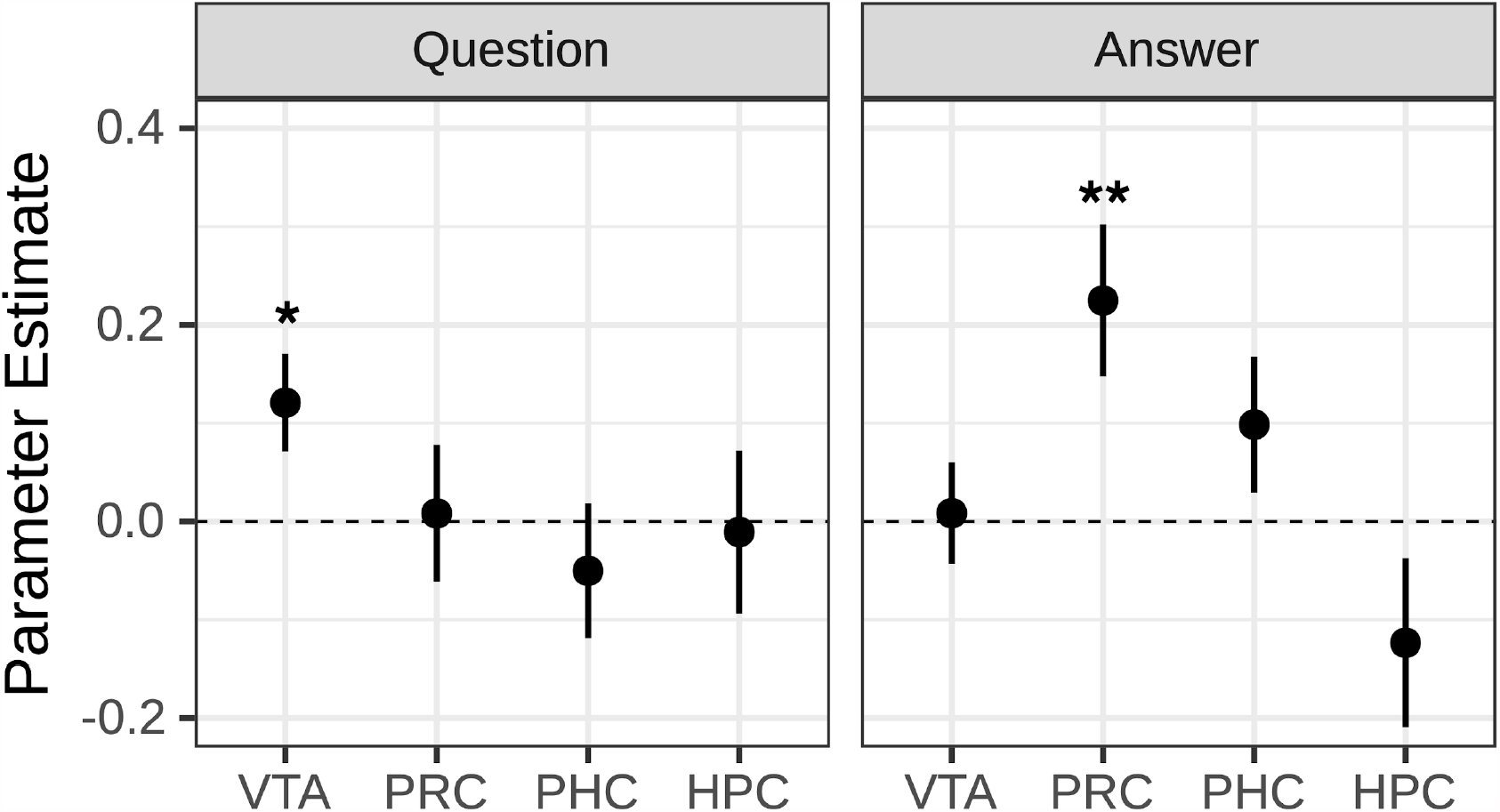
Parameter estimates for the logistic regression of memory recall with univariate activation across both the Question and Answer intervals. Anticipatory BOLD activation (during both the Question and Answer interval) in the VTA and medial temporal lobe ROIs was used to predict memory outcome for each trial in a mixed-effects logistic regression model. This method allows the identification of variance that is uniquely accounted for by each of the ROIs across both intervals. During the Question interval VTA activation was the only statistically significant predictor of subsequent recall of answers, while during the Answer interval PRC activation was the only statistically significant predictor of subsequent recall of answers. Error bars represent the SEM. ^**\***^ ***p*** < .05, ^**\*\***^ ***p*** < .01.

**Supplementary Figure 2.**
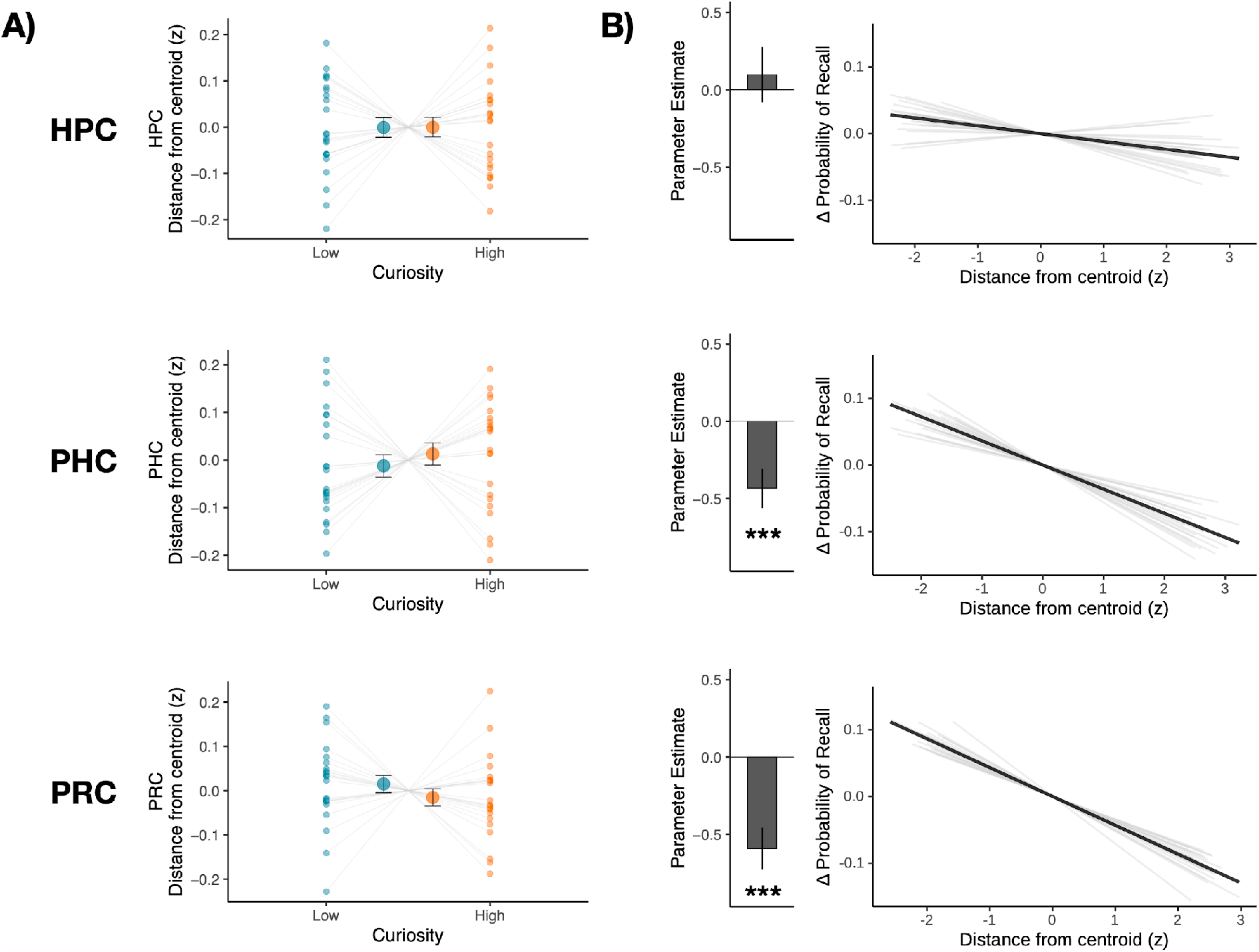
State convergence in the medial temporal lobe during the presentation of answers. **A)** During the presentation of trivia answers, curiosity did not influence convergence in any of the medial temporal lobe ROIs. The larger dots in each panel represents the group mean, while the smaller dots represent the mean distance for each participant. **B)** We used a mixed-effects logistic regression model to predict memory outcome for each trial using the state convergence of the medial temporal lobe ROIs. Convergence in the medial temporal cortices (PHC & PRC), but not the Hippocampus, were significant predictors of subsequent recall. Bar graph of each panel represents the parameter estimate of each ROI in the full model. For visualisation, the estimated change in probability of recall (demeaned within subject) is plotted against the distance from centroid for each ROI. Light gray lines depict the slope for each participant, while the solid black line depicts the mean slope across all participants. Error bars represent the SEM. ^***^ *p* < .001

